# Localization of Spatially Extended Brain Sources by Flexible Alternating Projection (Flex-AP)

**DOI:** 10.1101/2023.11.03.565461

**Authors:** Lukas Hecker, Amita Giri, Dimitrios Pantazis, Amir Adler

## Abstract

Magnetoencephalography (MEG) and electroencephalography (EEG) are widely employed techniques for the *in-vivo* measurement of neural activity with exceptional temporal resolution. Modeling the neural sources underlying these signals is of high interest for both neuroscience research and pathology. The method of Alternating Projection (AP) was recently shown to outperform the well-established recursively applied and projected multiple signal classification (RAP-MUSIC) algorithm. In this work, we further enhanced AP to allow for source extent estimation, a novel approach termed flexible extent AP (FLEX-AP). We found that FLEX-AP achieves significantly lower errors for spatially coherent sources compared to AP, RAP-MUSIC, and the corresponding extension, FLEX-RAP-MUSIC. We also found an advantage for discrete dipoles under forward modeling errors encountered in real-world scenarios. Together, our results indicate that the FLEX-AP method can unify dipole fitting and distributed source imaging into a single algorithm with promising accuracy.

## 1. INTRODUCTION

MEG and EEG (M/EEG) are crucial techniques for recording neural activity with high temporal precision. Despite their importance, a significant challenge in M/EEG is their low spatial resolution, complicating neural source localization due to an ill-posed inverse problem [1, 2].

Traditional localization approaches often rely on the least squares criterion over a source space, such as the minimum norm estimate (MNE) [3]. While widely used, MNE-based methods often struggle with identifying sparse neural sources. In contrast, dipole scanning techniques follow an alternate inversion strategy. Rather than estimating a distributed current density across the entire expansive source space, these methods focus on identifying a confined set of discrete sources. Typically, a localizer function is derived, and dipoles are shortlisted as candidates based on specific threshold criteria. A prominent technique in the M/EEG field is the recursively applied and projected Multiple Signal Classification (RAP-MUSIC, [4, 5]). In this iterative approach, individual dipoles are successively added to the set of active sources, and their signals are systematically removed from the measurements through an out-projection operation until a predefined criterion is met. A natural extension of this approach, the recent introduction of FLEX-RAP-MUSIC, allowed for the estimation of the spatial extent of neural sources [6]. The results showed that extent estimation is possible by smoothing the forward matrix with increasing order, thereby reducing localization errors significantly. A conceptually similar approach to RAP-MUSIC is Alternating Projection (AP) [7]. It differs from RAP-MUSIC in two key properties: (1) Unlike RAP-MUSIC, AP does not rely on the signal subspace but instead directly utilizes the data covariance. This property enhances the robustness of AP in localizing highly correlated sources, and (2) following the initial search, AP further refines the position of the discovered dipole through additional iterations. Despite being a widely adopted approach in sensor array signal processing, its advantage over many iterative scanning techniques was shown only recently [8]. Notably, AP exhibited significantly lower localization error compared to RAP-MUSIC in scenarios featuring highly correlated sources and low signal-to-noise ratios (SNR). This makes AP a viable choice in many real-world settings.

In this study, we combine the advantages of the AP approach and the FLEX framework to allow for the estimation of source extent (FLEX-AP). We compare FLEX-AP with other approaches and evaluate its efficacy under various conditions based on a wide variety of neural source simulations.

## 2. METHODS

### 2.1. Problem Formulation

M/EEG recordings ***y***(*t*) ∈ ℝ^*q*^, measured in *q* sensors at time *t*, can be modeled as the product of the forward matrix^1^ **G** ∈ ℝ^*q×p*^ and the current vector ***j***(*t*) ∈ ℝ^*p*^, where *p* is the number of candidate dipoles spanning the source space:

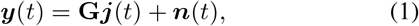

with ***n***(*t*) ∈ ℝ^*q*^ an additive noise term. The forward matrix describes the mapping of the sources of neural activity in the brain to the measurements recorded by the sensors, and takes into account the geometry of the head, the sensor positions, and the physical properties of the tissues between the sources and sensors. Note that, for simplicity, we focus here in the case of *fixed-orientated* dipoles, which are aligned with the aggregate direction of the dendrites of pyramidal cells, the primary generators of M/EEG signals [9, 10].

### 2.2. FLEX-AP

We describe here the calculation of the FLEX-AP inverse solution, which encompasses the standard AP approach as a special case with a maximum smoothness order of 0. In this work, we assumed that the number of active sources *Q* is known. To transition from a discrete dipoles to an extended source, we represent the mapping of a local patch of *coherent* brain activity to the sensors through an equivalent local smoothing operation on the forward matrix. Smoothness is achieved using a heat diffusion model, which simulates the concurrent activation of dipoles with diminishing amplitude as we move away from the center of the patch.

Mathematically, we first define the adjacency matrix **A** ∈ ℝ^*p×p*^ of the cortical mesh (triangulated surface) that spans the source space. The degree matrix **D** is then defined as the diagonal matrix with elements **D**_*ii*_ = Σ_*j*_ **A**_*ij*_. Using the Laplacian matrix **L** = **D** − **A** the diffusion operator **K** is computed as:

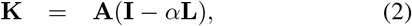

where 0 *< α <* 1 is a free diffusion parameter, and **I** ∈ ℝ^*p×p*^ is the identity matrix. With the diffusion operator **K**, we can generate a series of increasingly smoothed forward matrices to simulate sources of gradually greater extent, up to a maximum smoothness order of *k* :

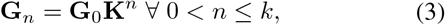

where **G**_0_ is the initial (“unsmoothed”) forward matrix and *n* reflects the order of the source extent, indicating the inclusion of neighbours up to the *n*-th order from the center of the patch.

Given this set of forward matrices, we then proceed with an exhaustive grid search over all candidate dipole locations *p* and source extents *k* to maximize the AP localizer function (equation 21 in [8]), yielding the initial (0-th) estimate of the first source:

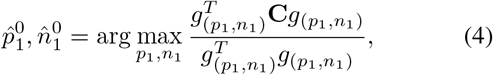

where **C** = **YY**^*T*^ ∈ ℝ^*q×q*^ is the data covariance matrix and *g*_(*p*_1_,*n*_1_)_ is the *p*_1_-th column of the forward matrix with the extent 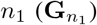. After detecting the initial source location and extent, we recursively add one source at a time, solving for the initial (0-th) estimate of the *q*-th source as:

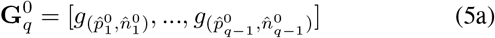

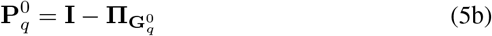

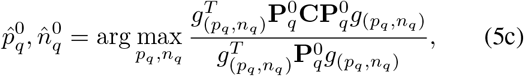

where 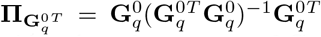 is the projection operator within the column space of **G**. Sources are detected recursively, *q* = 2, …, *Q*, until all sources *Q* are found. 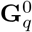 is the set of forward matrix columns corresponding to the previously selected source candidates. **P** denotes the projection matrix which projects out these components as part of the localizer function. This ensures that the next source candidate aligns with the data covariance matrix, after (2) the previously detected sources are projected out.

Once all the *Q* sources are found, the AP algorithm begins the refinement process of the location and extent of each sources over several iterations *j*:

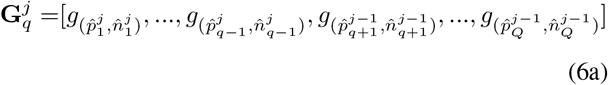

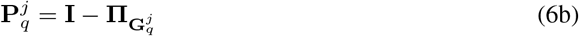

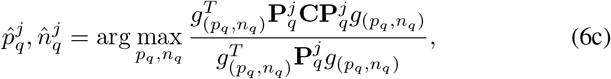

where *q* = 1, …, *Q* refines all sources. 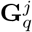 comprises the set of all but one forward matrix column, corresponding to the online estimate of the source candidates. The one not contained in the set is removed to find a potentially better candidate in the process. 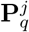 therefore out-projects all but one forward matrix column, and then the optimal new candidate is found through exhaustive search over all dipole locations and extents just as in Eq. (5). An illustration of the concept and impact of source extent estimation is shown in Fig. 1. We terminate the iterations of the AP algorithm after convergence, that is, when the location of the sources would not change [8].

**Fig. 1:**
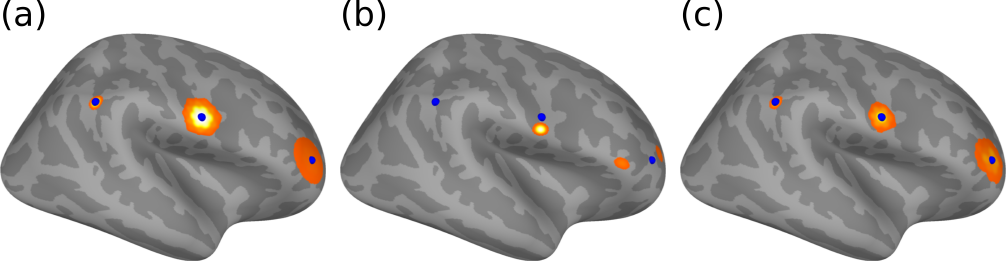
Illustration of Flexible Extent Estimation. (a) Simulation of a discrete source (left) and two increasingly extended sources (middle & right). (b) AP inverse solution results in biased (middle source) or multiple sources assigned to a single large extent source (right source). (c) FLEX-AP inverse solution yields an accurate solution. Blue spheres mark the center of the ground truth source locations depicted in (a).

The computational complexity of the FLEX-AP algorithm can be broken down to its four different stages, where (1) the covariance computation follows 𝒪 (*q*^3^), (2) the search for the initial source follows 𝒪 (*k* × *p* × *q*^2^), (3) the first round of iterations follows 𝒪 (*k* × *Q* × *q*^2^) and (4) the refinement stage follows 𝒪 (*k* × *Q* × *q*^3^). The dominant computational cost is tied to operations on the number of channels (*q*), but the most extensive interactions in terms of data size involve the number of dipoles (*p* ≫ *q*). Given the provided complexities and constraints, the algorithm is designed to handle larger number of dipoles effectively, while the maximum number of smoothness orders *k*, number of sources *Q*, and number of iterations have only a low impact on computational complexity. When comparing the FLEX-AP to the standard AP, the former exhibits a higher computational complexity that scales linearly with respect to the number of smoothness orders *k*.

### 2.3. Evaluation on Simulated EEG Data

For the evaluation we used a 128-channel EEG montage based on the international 10-20 system. The *fsaverage* template T1 image provided by the freesurfer software suite [11, 12] was used to model the cortical source space. As a base forward model, we used *p* = 5124 dipoles sampled along the cortical surface with fixed orientations. To avoid the pitfall known as the “inverse crime” [13], we employed two additional forward models to assess localization performance. First, to account for source space model errors, we used an alternate finer mesh comprising *p* = 8196 dipoles. Second, to further account for potential conductivity errors, we used the same *p* = 8196 dipole mesh while adjusting the skull-to-brain conductivity ratio from 1 : 80 to 1 : 50.

Every experimental condition included 500 Monte Carlo simulations, with each simulation considering two randomly located sources. To simulate the source time courses, we first generated two random sequences following a 1*/f* frequency spectrum. Subsequently, we generated the time courses of these two sources by mixing the sequences in a manner that adhered to a moderate inter-source correlation with a coefficient of *ρ* = 0.5 (as referenced in [14]). Within each condition, we examined these original sources with no extent (discrete source), but also with an extent of a given neighborhood order (extended source). To transform a discrete source into an extended source of order *n*, we applied the diffusion operator **K**^*n*^ to the source vector **j**(*t*), that is, **j**_ext_(*t*) = **K**^*n*^**j**(*t*). Last, we projected the sources to the sensors through the forward matrix, and added independent and identically-distributed noise to yield measurements with an SNR of 0dB.

We evaluated the FLEX-AP method against AP, RAP-MUSIC and FLEX-RAP-MUSIC. Note, the smoothing of forward matrices in the FLEX-RAP-MUSIC algorithm as described in [6] was corrected to follow the heat diffusion model. Thus, both FLEX-AP and FLEX-RAP-MUSIC employed an identical set of smoothed forward models (Eq. (3)). For the evaluation, we used two evaluation metrics: (1) Mean Localization Error (MLE), calculated as proposed by [15]. For extended sources, we identified the maxima of the source vector prior to the MLE calculation. (2) Earth Mover’s Distance (EMD), a measure that takes into account the error in extent estimation, as introduced in [16]. For the calculation of EMD, we used the python package *POT* [17].

## 3. PERFORMANCE EVALUATION

FLEX-AP was developed with the capacity to localize both discrete and extended sources. Figure 2 shows the MLE and EMD for all methods with sources of increasing source extent. A neighborhood order of 0 indicates discrete sources, while an order of 8 suggests a large extended source of significant size.

**Fig. 2:**
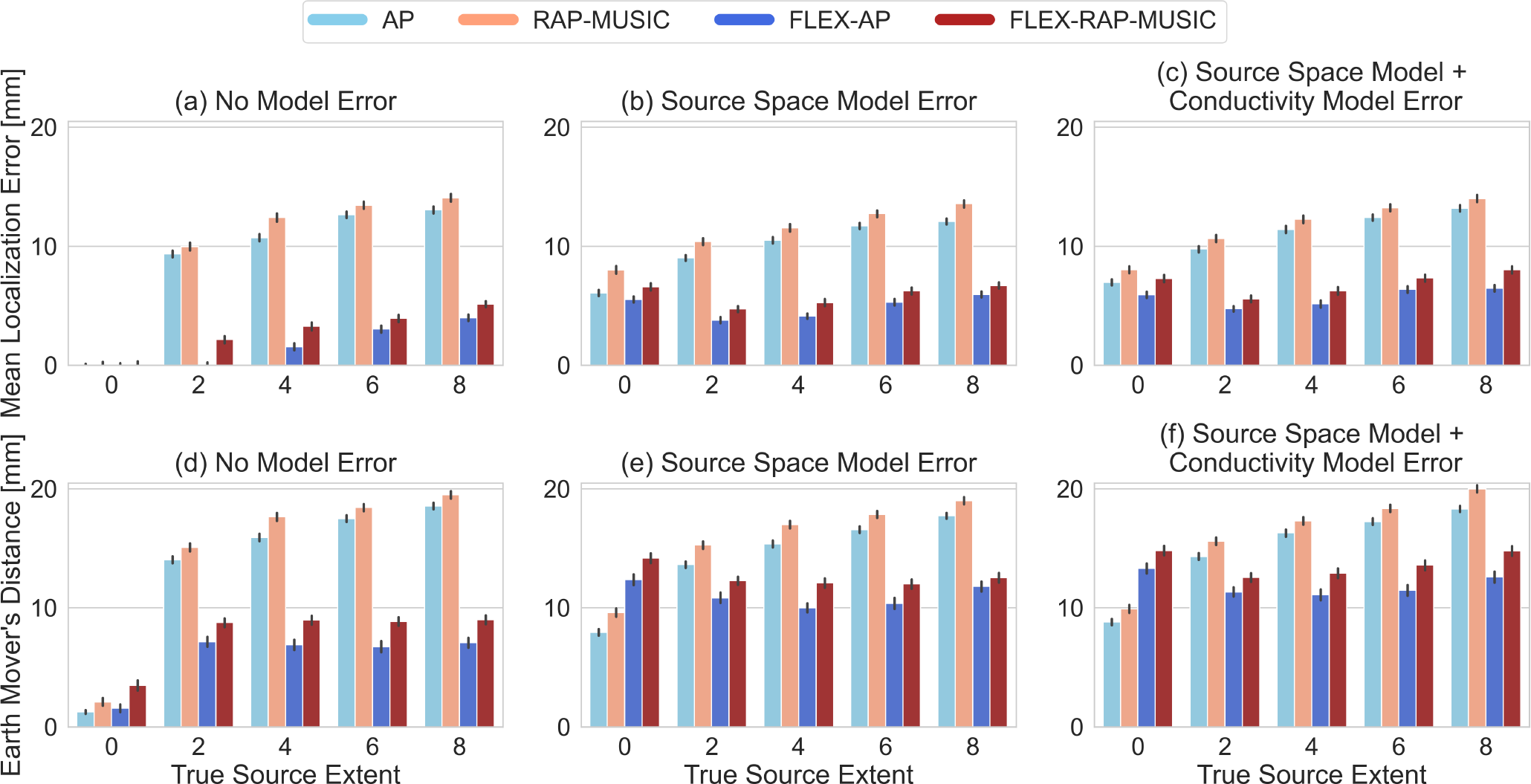
Localization Performance for Different Source Extents: (a)-(c) Mean Localization Error; and (d)-(f) Earth Mover’s Distance. The left column represents inversion with the same forward model as used for data generation. The center column shows inversion on a different source space grid and the right column shows inversion on a different grid and a mismatch of skull-to-brain conductivity ratio between simulation and inversion. Source extent is denoted by neighborhood orders, where 0 corresponds to a discrete source. A neighborhood order of 2, 4, 6, and 8 corresponds to an average active cortical surface area of 0.26 cm^2^, 1.11 cm^2^, 2.51 cm^2^, and 4.46 cm^2^, respectively. Bars show the median error over 500 Monte Carlo simulations.

First, we highlight that our results replicated our prior work introducing AP for the M/EEG inverse problem [8]. In our experiments, AP consistently outperformed RAP-MUSIC in terms of localization errors, as measured by both MLE and EMD.

In the absence of forward modeling error (Fig. 2(a & d)), we observed that AP only exhibited a small advantage over the FLEX-AP algorithm when the true source extent was 0. However, as source extent increased, both FLEX-AP and FLEX-RAP-MUSIC demonstrated significantly lower errors compared to their non-FLEX counterparts, both in terms of MLE and EMD. This suggests that accounting for spatial extent estimation improved localization. Similar to the comparison between AP and RAP-MUSIC, FLEX-AP consistently outperformed FLEX-RAP-MUSIC across all scenarios presented.

Remarkably, in the presence of source modeling error, we found that the FLEX methods exhibited lower MLE for discrete sources than their non-FLEX analogues (Fig. 2bc). Figure 3 illustrates this behavior with two examples from our simulations. Notably, the extended sources identified by FLEX-AP closely aligned with the ground truth positions, while the discrete sources located by AP were displaced from their true positions.

**Fig. 3:**
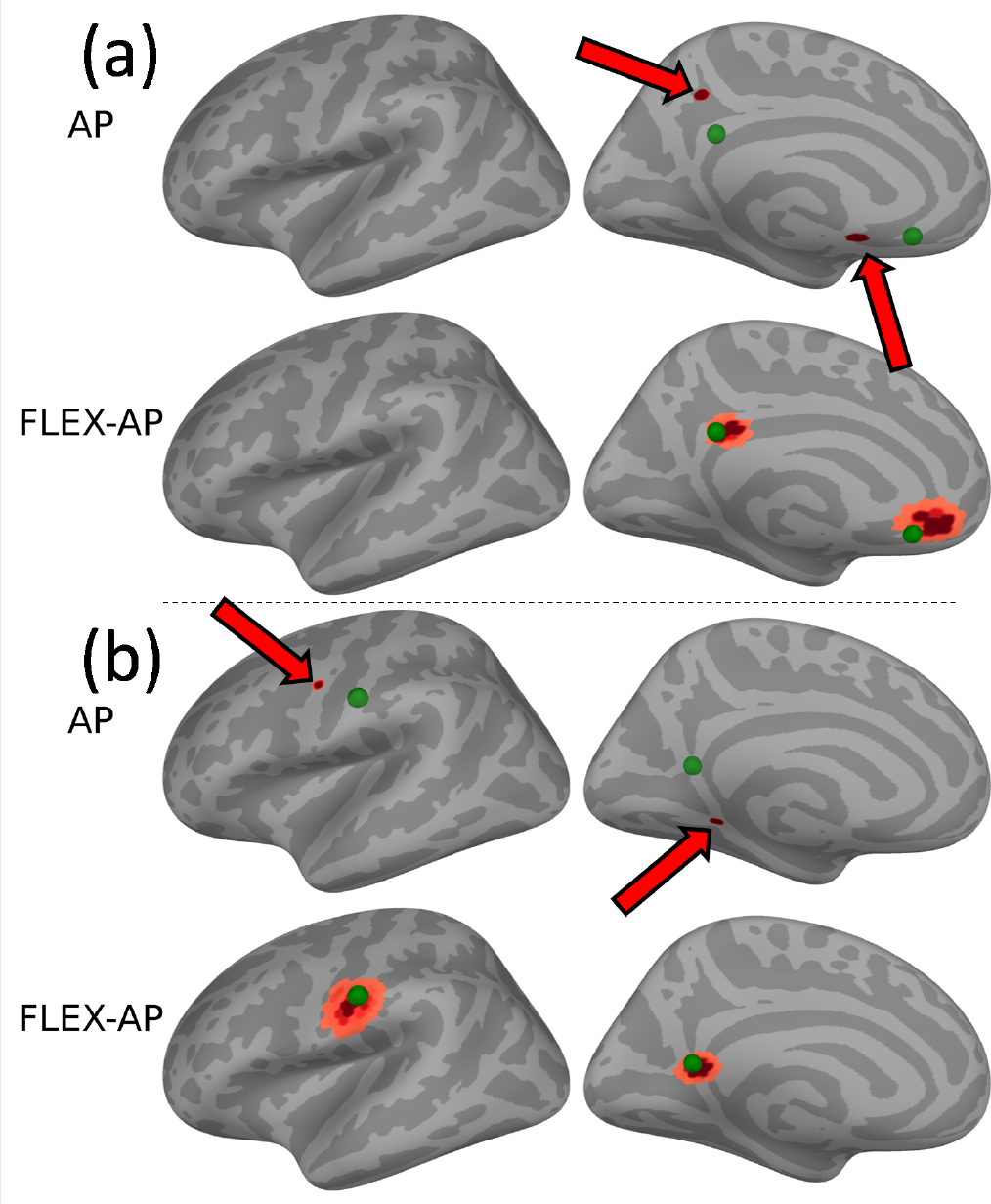
Localization of Discrete Sources under Source Space Modeling Error. Two instances showcasing pairs of simulated discrete sources, depicted as green spheres (a and b). The red patches indicate the inverse solution. Notably, the AP solutions are displaced, whereas the sources found using FLEX-AP are centered correctly.

## 4. CONCLUSIONS

We introduced an extension to the AP algorithm that enables flexible extent estimation for neural sources. Our results indicate that incorporating extent estimation leads to a reduction in overall localization errors. Importantly, we also found that extent estimation can mitigate localization errors, even when dealing with discrete sources amidst source modeling errors, a scenario commonly encountered in real-world data. We further validated that AP framework is superior to the more widely adopted RAP-MUSIC algorithm.

Future research is needed to validate these findings on real data, and expand the analyses in uncorrelated sources. Furthermore, the diffusion parameter *α* (cf. Eq. 2) carries assumptions about the distribution of currents within an extended source. Future investigations could study the influence of this parameter and consider the potential expansion of the FLEX-AP algorithm to accommodate sources with varying degrees of diffusion, and even scenarios where current distributions may deviate from typical center-focused patterns.

## 5. DATA & CODE AVAILABILITY

Open source code to reproduce these results is available at: https://github.com/LukeTheHecker/ap_paper_analyses. A python library implementing FLEX-AP together with other inverse solutions is available at: https://github.com/LukeTheHecker/invert. An implementation of AP in MATLAB can be found at https://alternatingprojection.github.io/.

## 6. COMPLIANCE WITH ETHICAL STANDARDS

This is a numerical simulation study for which *no* ethical approval was required.

## 7. ACKNOWLEDGEMENTS

This work was supported by the United States-Israel Binational Science Foundation grant 2020805 to A.A. and NIH grant 1R01EY033638 to D.P.. The authors have no relevant personal financial or non-financial interests to disclose.

## Footnotes

1 The forward matrix is also referred to as *lead field* or *gain matrix*.

